# Inconsistent patterns of microbial diversity and composition between highly similar sequencing protocols: a case study with reef-building corals

**DOI:** 10.1101/2021.06.30.450656

**Authors:** Hannah E. Epstein, Alejandra Hernandez-Agreda, Samuel Starko, Julia K. Baum, Rebecca Vega Thurber

## Abstract

16S rRNA gene profiling (amplicon sequencing) is a popular technique for understanding host-associated and environmental microbial communities. Most protocols for sequencing amplicon libraries follow a standardized pipeline that can differ slightly depending on laboratory facility and user. Given that the same variable region of the 16S gene is targeted, it is generally accepted that sequencing output from differing protocols are comparable and this assumption underlies our ability to identify universal patterns in microbial dynamics through meta-analyses. However, discrepant results from a combined 16S rRNA dataset prepared by two labs whose protocols differed only in DNA polymerase and sequencing platform led us to scrutinize the outputs and challenge the idea of confidently combining them for standard microbiome analysis. Using technical replicates of reef-building coral samples from two species, *Montipora aequituberculata* and *Porites lobata*, we evaluated the consistency of alpha and beta diversity metrics between data resulting from these highly similar protocols. While we found minimal variation in alpha diversity between platform, significant differences were revealed with most beta diversity metrics, dependent on host species. These inconsistencies persisted following removal of low abundance taxa and when comparing across higher taxonomic levels, suggesting that bacterial community differences associated with sequencing protocol are likely to be context dependent and difficult to correct without extensive validation work. The results of this study encourage caution in the statistical comparison and interpretation of studies that combine rRNA sequence data from distinct protocols and point to a need for further work identifying mechanistic causes of these observed differences.

**Importance:** Amplicon sequencing remains a popular technique for characterizing organism and environmental microbiomes. The publication of sequence data from microbiome studies on open-access repositories provides an opportunity to identify universal patterns in microbial dynamics. To this end, it has been widely accepted that sequencing output from differing protocols are comparable and can be combined for analysis, so long as the same gene region is targeted. While most protocols for amplicon sequencing follow standardized pipelines, they can differ slightly between laboratory facility and user. In this study, we compared technical replicates of coral samples to evaluate the efficacy of combining organism-associated microbial datasets derived from two differing protocols. We found inconsistencies in the differences between bacterial communities, which persisted following data manipulations intended to increase comparability. These results suggest caution must be taken in the statistical comparison and interpretation of studies that combine data derived from distinct protocols.

## Introduction

Microbial ecology has benefited tremendously from recent technological advances in areas such as high throughput sequencing [1]. The generation of large volumes of genomic data (e.g., 16S rRNA) has encouraged large-scale collaborative efforts, including the Human Microbiome Project (HMP; https://www.hmpdacc.org/hmp [2]), the Earth Microbiome Project (EMP; www.earthmicrobiomeproject.org [3]) and TARA Oceans [4], which aim to catalogue all microbial life associated with humans, other animal hosts and across ecosystems. Publicly available sequencing data resulting from these initiatives provide opportunities for the production of meta-analyses and for researchers with smaller scale projects to make comparisons and/or combine their dataset with a much broader set of samples, allowing increased impact of their finer sequencing efforts.

Large-scale collaborations also provide standardized protocols for replication and adequate comparison. For example, the EMP standardized protocols for 16S rRNA sequencing are optimised for repeatedly processing large numbers of samples and benefit from automation and high throughput sequencing on an Illumina HiSeq platform. Smaller, individual research laboratories are, in many cases, processing fewer samples less frequently, likely without access to automation, but with the capacity to shift reagents and polymerase chain reaction (PCR) conditions to achieve optimised results. Smaller numbers of samples are also more often sequenced on the Illumina MiSeq platform due to cost effectiveness and increased read length. It has been previously accepted that HiSeq and MiSeq platforms produce comparable results (see [5]). In fact, there are few differences in the two sequencing platforms: apart from the discrepancy in read length (HiSeq: 150bp; MiSeq: up to 300bp) and sequencing depth (HiSeq: 150M reads/lane; MiSeq: 20-25M reads/lane), the chemistry between the two methods is almost identical, except for the slightly different concentrations of sodium hydroxide (NaOH) used to denature the libraries for sequencing (HiSeq: 0.1N NaOH; MiSeq: 0.2N NaOH; outlined in [6]). As a result, meta-analyses of 16S rRNA data across microbial study systems already utilize cross-protocol and platform data that are stored in public repositories (see [7–9]).

However, when attempting to combine 16S rRNA data for a large, longitudinal coral microbiome dataset, we found that the data derived from our in-house preparation and MiSeq sequencing runs clustered separately from those prepared and sequenced by EMP, despite following a highly similar preparation protocol. This led us to re-evaluate if the two protocols utilising different sequencing platforms provide comparable results. Using 24 coral samples that were sequenced in parallel both in-house (MiSeq) and by EMP (HiSeq), we examined if methodological biases lie within these complex microbial communities, and how (or whether) results obtained from the two protocols are comparable when running standard microbial ecology analyses on alpha diversity, beta diversity, dispersion and differential abundances. Large collaborative sequencing efforts and public sharing of these data are central to understanding general, cosmopolitan patterns in the coral microbiome, which makes effective comparison of sequencing data originating from multiple laboratories vital.

## Methods

### Sample collection, DNA extraction, library preparation and sequencing

Coral samples were originally collected from Kiritimati (Christmas) Island in May 2015 from two species: *Porites lobata* (n = 13) and *Montipora aequituberculata* (n= 11). Frozen tissue was sent directly to EMP (University of California, San Diego) for DNA extraction, PCR, library preparation and sequencing on an Illumina HiSeq 2×150bp run [10]. Frozen tissue from the same samples were also processed in-house at Oregon State University (see previously published methods in [11]) using a highly similar protocol as EMP but sequenced on an Illumina MiSeq 2×300bp run. Both protocols targeted the V4 region of the 16S rRNA gene with the following primers: 515F [12] 5’–<u>TCGTCGGCAGCGTCAGATGTGTATAAGAGACAG</u>GTGYCAGCMGCCGCGGTAA–3’ and 806R [13] 5’–<u>GTCTCGTGGGCTCGGAGATGTGTATAAGAGACAG</u>GGACTACNVGGGTWTCTAAT-3’, with the Illumina adapter overhangs underlined. The only difference between the two protocols was the Taq used for PCR: EMP used Platinum Hot Start PCR MasterMix (Thermofisher) and in-house used Accustart^™^ II PCR ToughMix (QuantaBio). Hereafter, the two protocols will be referred to by their most significant difference: the “HiSeq protocol” run by EMP and the “MiSeq protocol” run in-house.

### Bioinformatics

Sequences from both HiSeq and MiSeq protocol outputs were combined into a single dataset and processed using the QIIME2 pipeline to undergo trimming, quality control, identification of amplicon sequence variants (ASVs), and taxonomic assignment. To ensure comparability between the two protocols and accuracy in ASV-picking, we chose to follow similar treatment of sequencing data by EMP [3]; only forward reads were used and trimmed to 120bp. Primers were removed using the plug-in cutadapt [14], and denoising and ASV picking was performed using the DADA2 plug-in [15]. For comparison, ASVs were simultaneously clustered using the plugin vsearch [16] by 97% similarity, resulting in two output biom tables: one for ASVs and one for 97% clustered operational taxonomic units (hereafter, “OTUs”). Taxonomic assignment for both tables was performed using a naïve Bayes classifier with the SILVA v. 132 database [17], trained on each set of representative sequences from the two pipelines.

### Data import into R

All statistical analyses were performed in R v. 4.0.2 [18]; graphics were conducted in R using the package ggplot2 [19]. QIIME feature tables, taxonomic assignments, and tree files for the ASV and OTU datasets were imported into phyloseq [20] via qiime2R [21] for downstream analyses. The SILVA annotations characterized some reads as Phylum: Alphaproteobacteria, Family: Mitochondria. This annotated family contained a mix of bacterial and mitochondrial (eukaryotic) reads: thus eukaryotic mitochondria were further identified using BLASTn [22] and subsequently removed from the two datasets. In the absence of blank controls from the EMP dataset, contaminants were identified using the 4 blank control samples from the MiSeq Data [11]. Contaminants were identified and removed (n=102) by prevalence using the decontam package [23] with a threshold value of 0.5 to ensure all sequences that were more prevalent in negative controls than positive samples were removed. Samples with less than 1000 and 998 reads, respectively, were removed from all analyses for ASV and OTU data. These two numbers differ slightly due to differences in contaminant and mitochondrial read removals as a result of ASV identification versus OTU picking.

### Diversity metrics and differential abundance

We conducted all diversity tests on the two coral species separately due to well-established differences in both alpha and beta diversity measures across host species [24–26] that could have obscured significant differences between protocols. Three alpha diversity metrics were calculated to account for richness, evenness and phylogenetic diversity: observed species richness, Shannon diversity index and Faith’s Phylogenetic Diversity (PD) were calculated on rarefied data (1000 and 998 reads/sample for ASV and OTU data, respectively). For each host species and data type (ASV and OTU), the three alpha diversity metrics were checked for normality using standardized residual plots, Q-Q plots and Shapiro-Wilk tests. If required, log and square-root transformations were performed to meet assumptions (see Table S1). Differences in alpha diversity indices between protocols were tested using paired t-tests. We also quantified four metrics of beta diversity to examine differences in microbial communities accounting for microbial abundance (Bray Curtis), presence/absence (binary Jaccard), phylogeny coupled with abundance (weighted Unifrac) and phylogeny coupled with presence/absence (unweighted Unifrac). For each host species, we constructed Bray-Curtis and weighted Unifrac dissimilarity matrices using the relative abundances of taxa to account for differences in sequencing depth between data derived from HiSeq and MiSeq protocols and constructed binary Jaccard and unweighted Unifrac dissimilarity using unrarefied counts. Dissimilarity matrices for all metrics were also built with unrarefied data after removing rare taxa (abundance below 0.5% and 1% threshold). Differences in beta diversity (i.e. both multivariate location (‘turnover’ and variation) were tested using permutational analyses of variance (PERMANOVAs) with 999 permutations blocked by coral colony ID (strata = sample label) using the adonis function from the package vegan [27] implemented in phyloseq [20]. Homogeneity of variances were further tested between protocols using betadisper with 999 permutations and communities were visualised using non-metric multidimensional scaling (NMDS) plots. To identify specific significant differences in taxon abundance in the two protocols, differential abundance analyses were performed using DESeq2 [28] on unrarefied count data with an alpha cut-off of 0.05. All analyses were performed on both ASV and OTU datasets unless otherwise specified. To assess any differences in secondary structure between specific ASVs differentially abundant in HiSeq vs. MiSeq protocols resulting from library denaturation, GC content (%) and melting temperatures were verified through the TmCalculator package in R [29]. Differences in mean GC content and melting temperature were tested among ASVs present in MiSeq, HiSeq and both protocols using analyses of variance (ANOVAs). We chose to present results from unrarefied data unless otherwise specified in the above methods; however, all analyses were run on both rarefied and unrarefied data and showed no major differences in significance (see Supplementary Tables S2, S4 & S5).

## Results

### Sequencing results

To test whether coral microbiome sequence data generated from the two protocols were comparable, we analysed paired sequence libraries, combined for comparative downstream analyses, using four standard variables for assessing microbiome variations at both the ASV and OTU levels: alpha diversity, beta diversity, beta dispersion, and differential abundance measures. The final ASV dataset included all 24 samples (total n = 48 to account for 2 technical replicates per sample, one from each protocol) with a combined total of 1,444,493 reads consisting of 5,512 distinct ASVs for analysis. In the OTU dataset, two *P. lobata* HiSeq protocol samples contained less than 998 reads and were removed along with their MiSeq protocol counterparts leaving 22 samples (total n = 44 to account for 2 technical replicates per sample, one from each protocol) for comparison consisting of 953,396 reads and 2,174 OTUs. All comparisons were done using identical read values for both protocols.

### Sensitivity of alpha diversity to protocol

When using ASVs for analysis, there was a slight tendency for alpha diversity to be lower when calculated from MiSeq protocol data, but diversity did not differ significantly between protocols for any of the four alpha diversity metrics measured (*p* > 0.05: Table S1), in either species (Figure 1). However, when using OTUs there was a host species-specific effect on some measures of diversity. Specifically, alpha diversity was not significantly different between protocols for *M. aequituberculata*, but there was protocol sensitivity for *P. lobata* when using Shannon diversity and Faith’s PD measures (but not observed richness), both of which were significantly greater in data resulting from the HiSeq protocol (Figure S1; Table S1).

**Figure 1:**
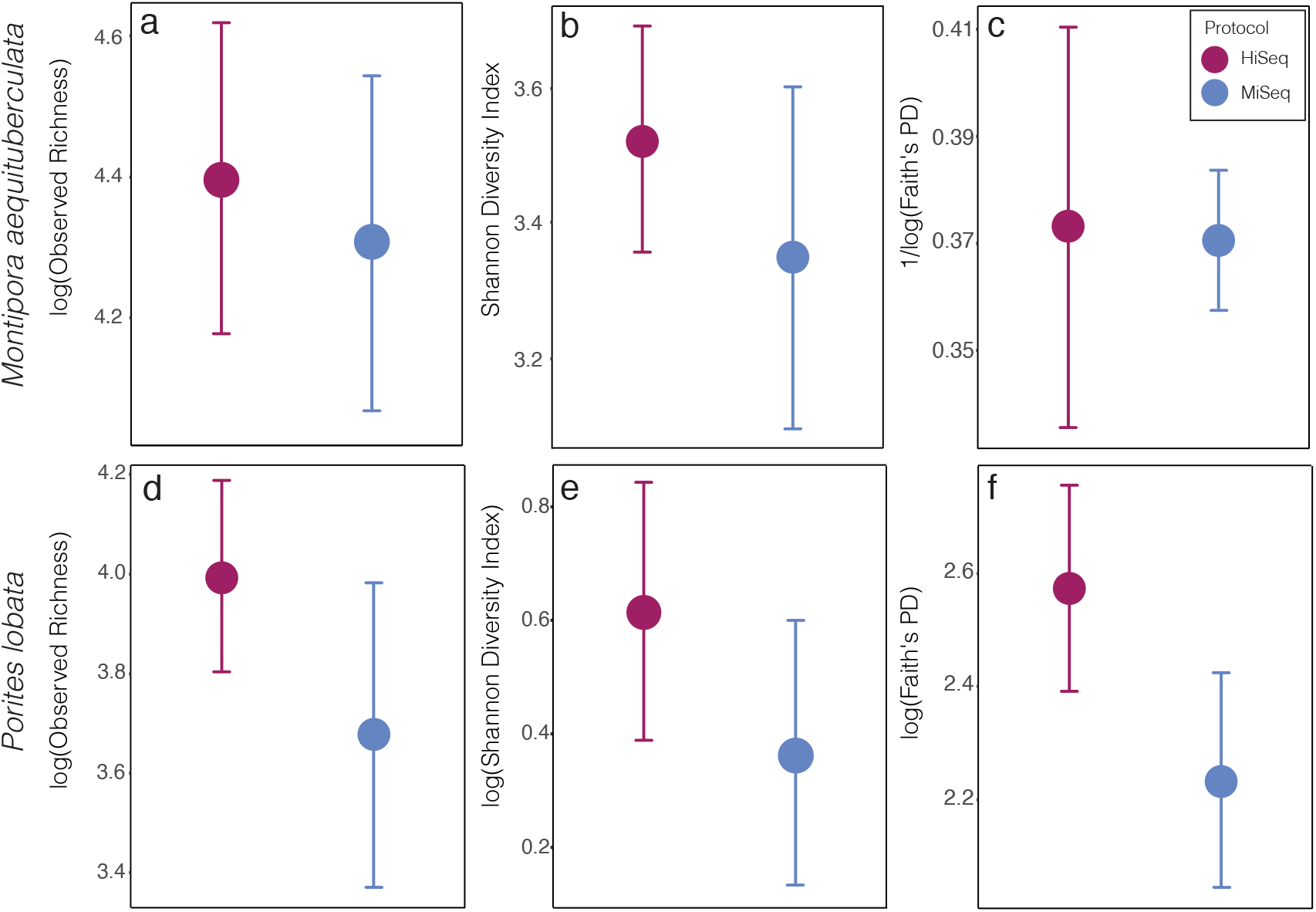
Alpha diversity metrics of bacterial ASVs between protocols in both species, *Montipora aequituberculata* (a-c) and *Porites lobata* (d-f), for observed species richness (a, d), Shannon diversity (b, e), and Faith’s phylogenetic distance (c, f) (p>0.05 for all tests, see Table S1).

### Protocol explains large amount of variation in community beta-diversity

Significant differences in microbial community composition were found between protocols for both *M. aequituberculata* and *P. lobata* in all beta diversity metrics except for weighted UniFrac distances for *M. aequituberculata* in both ASV and OTU datasets (Figures 2 & S2; Table S2). All beta diversity metrics maintained similar dispersions (homogeneity of variances), aside from Bray Curtis for *P. lobata* in both ASV and OTU datasets (Figures 2g & S2g), as well as Unweighted UniFrac distances for *M. aequituburculata* in the ASV dataset (Figure 2b) and *P. lobata* in the OTU dataset (Figure S2f) (Table S2). While the communities did not show consistent, distinct visual segregation of nMDS data clouds according to protocol (Figures 2 & S2), some individual samples had highly different relative abundances of bacterial taxa (Figures 3). While the top 10 most abundant taxa were similar between protocols and across datasets, differences in the relative abundances and detection of some phyla were present (Figure 3; Table S3). In the ASV dataset, seven out of ten phyla were detected in both MiSeq and HiSeq protocols, and phylum-level bacterial community compositions across samples were dominated by Proteobacteria, followed by Firmicutes. However, when clustered as ASVs, these two phyla account for 74.71% versus 54.67% of the composition in MiSeq and HiSeq protocols, respectively, and three different phyla were alternatively detected between the platforms. One of them was phyla Euryarchaeota, which was present in the ASV dataset for MiSeq protocol samples with a mean relative abundance of 4.45% (Table S3), but absent in the top 10 most abundant taxa for HiSeq protocol samples, in which the mean relative abundance was less than 0.002%. Although differences in the relative abundances were persistent when clustering at the 97% OTU level, fewer discrepancies were observed (Figure 3; Table S3). For example, nine out of ten phyla were detected in both protocols, and the two dominant phyla (Proteobacteria and Firmicutes) account for 79.1% and 75.15% for MiSeq and HiSeq protocols, respectively. Interestingly, in both ASV and OTU datasets, the most abundant phyla were more evenly represented across samples from the HiSeq protocol as opposed to the MiSeq protocol (see “n” in Table S3). However, libraries derived from the HiSeq protocol and, to a lesser extent the MiSeq protocol, also contained several unclassified bacterial ASVs that were resolved when clustering at the 97% OTU level (see “NA” in Figure 3), suggesting some challenges in the taxonomic assignment of ASVs from short read HiSeq data.

**Figure 2.**
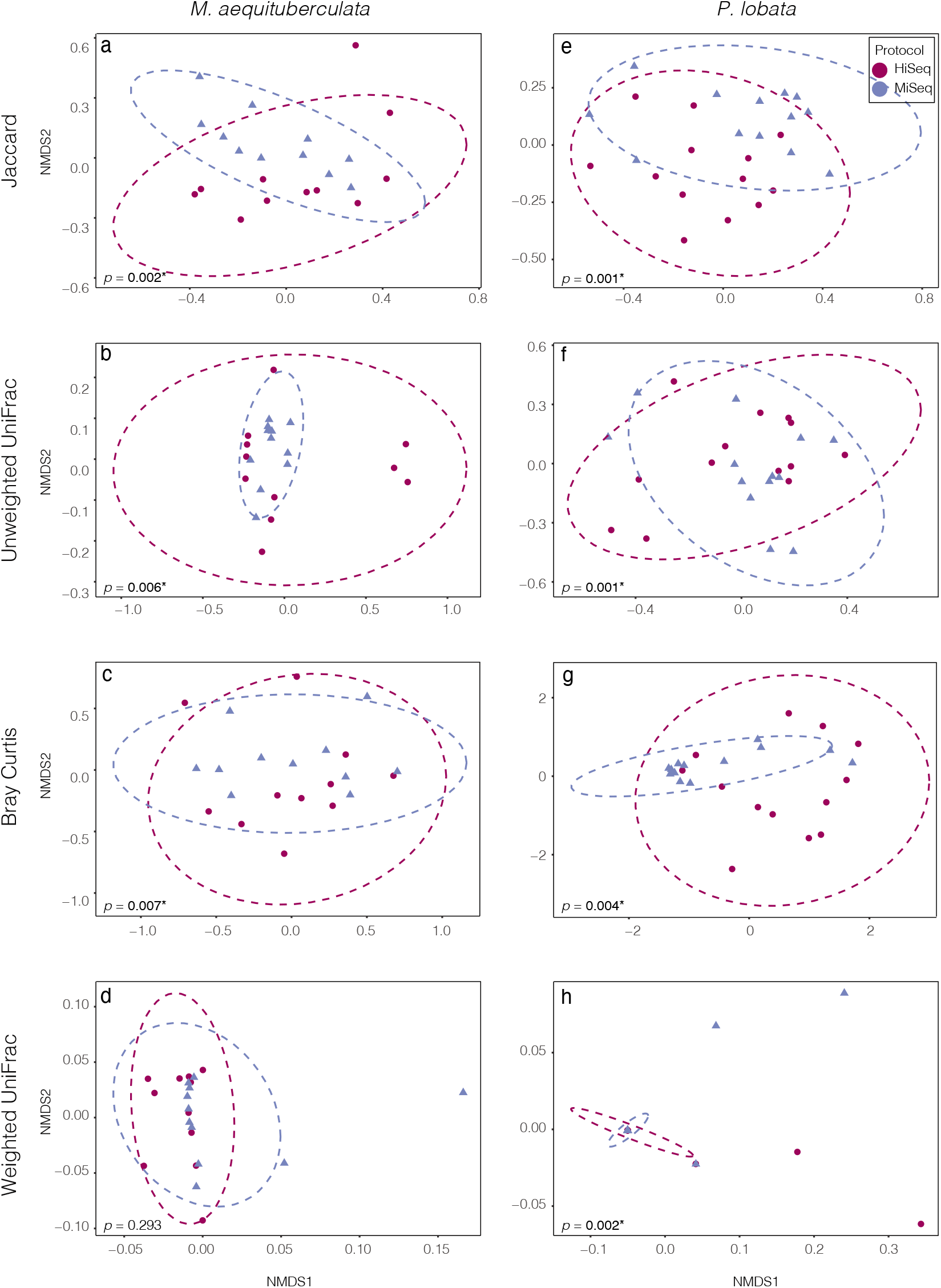
Non-metric multidimensional scaling (NMDS) ordinations of ASV bacterial communities between the two species, *M. aequituberculata* (a-d) and *P. lobata* (e-h) for each of the four tested dissimilarity metrics: Jaccard, Unweighted UniFrac, Bray Curtis and Weighted UniFrac. *P*-values with asterisks (*) refer to significant PERMANOVA results (see Table S2).

**Figure 3.**
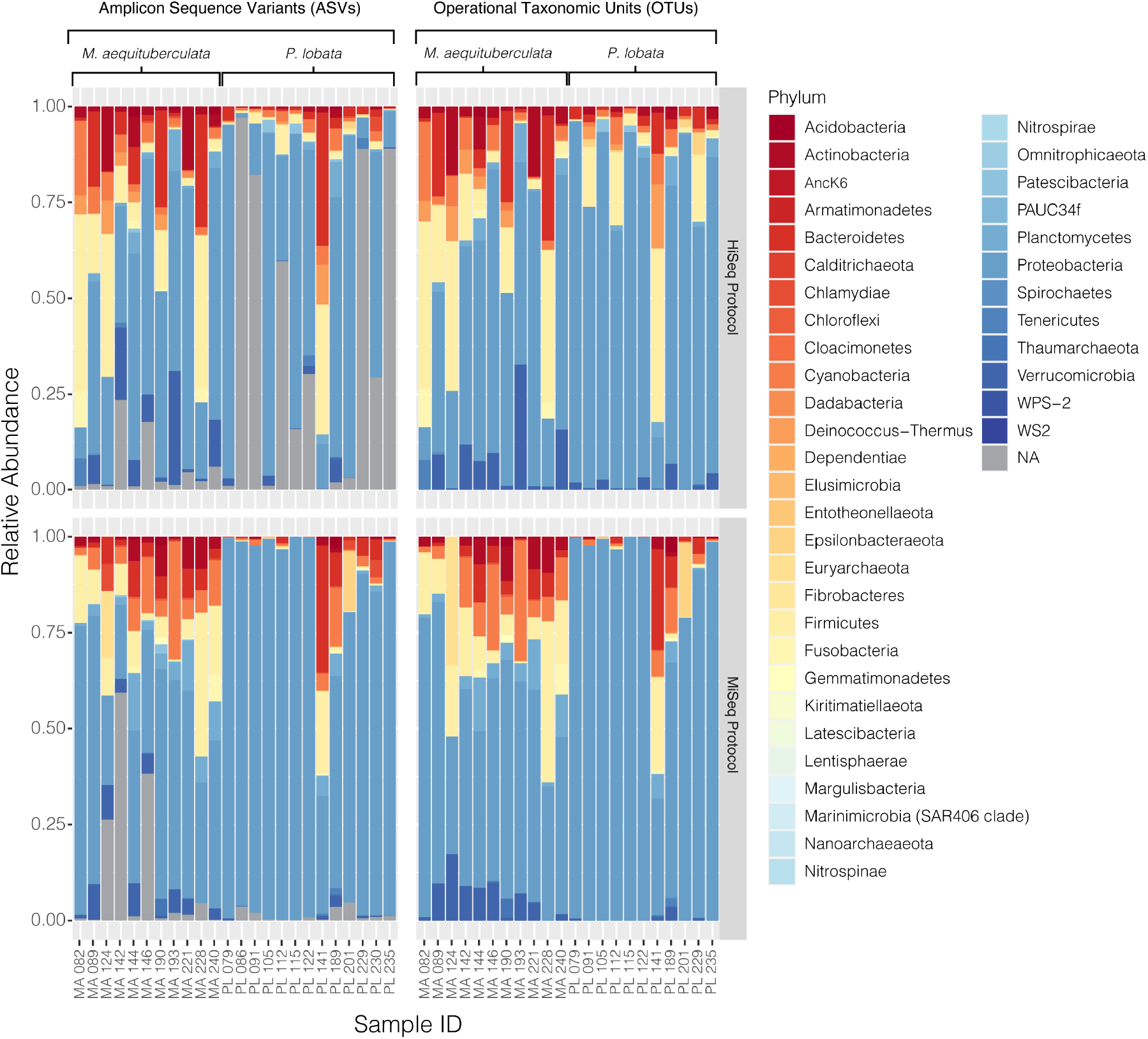
Relative abundances of bacterial phyla present in each coral sample from both *M. aequituberculata* and *P. lobata* prepared and sequenced using the HiSeq protocol (top) and the MiSeq protocol(bottom), using both the ASV (left) and OTU (right) datasets. “NA” refers to unassigned bacterial reads.

### Standard Normalizations Do Not Overcome Protocol Induced Variability in Microbiome Diversity

To examine whether standard normalization methods used in the field could overcome the differences between protocols, the datasets were manipulated by either removing rare taxa or grouping at higher taxonomic classifications, including Family and Phylum level. Truncating the microbial communities by removing rare taxa did not eliminate the beta diversity differences between the two protocols (Table S4). Removing rare taxa reduced the communities to less than 20 taxa (at ASV or OTU level), representing less than 1% of the total and suggesting that these bacterial communities are predominantly composed of low abundance taxa. Datasets of both species and taxonomic assignments (ASVs and OTUs) maintained the previously seen significant differences between protocols for all dissimilarity metrics aside from Weighted Unifrac for *M. aequituberculata* for both 0.5% and 1% rare taxa cut-offs (Table S4). To reduce the effects of minor differences in closely related bacterial taxa, we also ran PERMANOVAs and homogeneity of variance tests on communities at both the Family and Phylum classification level. Significant differences were found again between protocols, however this varied according to both host species and taxonomic level (Table S5). *Porites lobata* showed significant differences between protocols even at the Phylum level, whereas *M. aequituberculata* communities were significantly different between protocols at the Family level, but only the two dissimilarity metrics utilising presence/absence data (binary Jaccard and Unweighted UniFrac) showed significant differences at the Phylum level.

### Differential Abundance Analysis is Not Protocol Agnostic

Differential abundance analyses showed that only a few specific ASVs were significantly enriched in one protocol or the other (Figure 4; Table S6). The most enriched taxa belong to the dominant phyla, Proteobacteria and Firmicutes, with the magnitude of enrichments ranging between an approximately 7- and 29-fold change. While there was variation in differentially abundant taxa between protocols according to species and clustering method, some taxa were consistently different. For instance, HiSeq protocol libraries for both species had consistently higher abundances of *Geobacillus* sp., and lower abundances of Xenococcus PCC-7305 using the OTU dataset (Figure 4b & d). The magnitude of these enrichments was also consistent between coral species (Table S6). Importantly, differential enrichment between the protocols was observed in two taxa identified as crucial players of coral health and resilience, *Endozoicomonas* and *Vibrio* spp. *Endozoicomonas* exhibited significantly higher abundances in data derived from the HiSeq protocol in both species and datasets (ASV vs OTU), except in *P. lobata* using the OTU dataset (Figure 4a-c). *Porites lobata* showed significantly higher abundances of a *Vibrio* ASV when sequences were prepared with the MiSeq protocol (Figure 4c), but this difference was not maintained in the OTU dataset (Figure 4d). A closer look at all *Vibrio* and *Endozoicomonas* ASVs found in the sequencing output from HiSeq, MiSeq, or both protocols revealed no significant differences in mean GC content or melting temperatures (Table S7).

**Figure 4.**
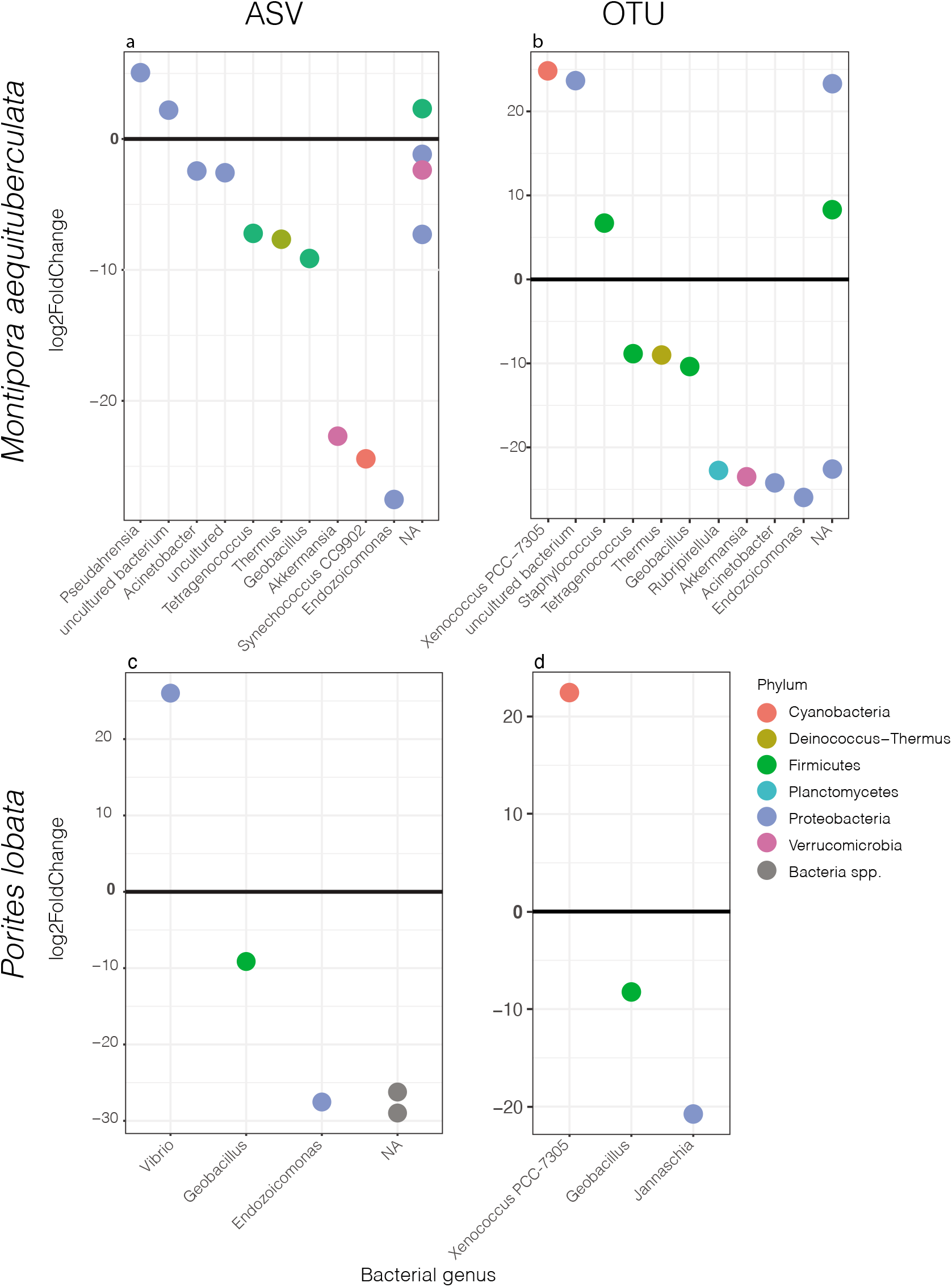
Significantly different ASVs (left) and OTUs (right) between protocols labelled by bacterial genus and coloured by phylum for *M. aequituberculata* (top) and *P. lobata* (bottom). Positive log2fold change refers to those significantly enriched in MiSeq protocol samples and negative log2fold change are those significantly enriched in HiSeq protocol samples. NA refers to bacteria unclassified at genus level.

## Discussion

In Scleractinian corals, 16S rRNA gene profiling remains a common and cost-effective tool for quantifying diversity of bacteria and some archaea in holobionts [30]. With the increase in sequencing of coral hosts by large collaborative groups such as EMP, and subsequent public sharing of sequencing data, it has become a common goal to examine widespread patterns through meta-analyses that combine datasets from multiple laboratories, making direct comparability a necessity.

In this small-scale comparative analysis of technical replicates, we found the greatest differences between the HiSeq and MiSeq protocols in the beta diversity and dispersion measures of coral microbiomes. Specifically, beta diversity and dispersion metrics were inconsistent between protocols, host species, and dissimilarity metrics, with differences in protocol explaining between 4 - 28% of the microbiome variability. Certain taxa were also significantly enriched in only one of the protocols, including those with known ecological importance. For example, *Vibrio* spp. and *Endozoicomonas* spp. ASVs were significantly enriched in MiSeq and HiSeq protocols, respectively. These two taxa have been identified as important in the health and maintenance of coral homeostasis and are often used to make statements about the health of the coral host [31, 32]: *Vibrio* spp. have been implicated in disease [33] but remain common partners in healthy corals, while *Endozoicomonas* spp. are hypothesized to benefit to coral health via synthesis of dimethylsulfoniopropionate (DMSP) [34], carbohydrate cycling and protein provisioning [35], and may be considered a potential symbiont [36]. The differential abundances of these two taxa between protocols are particularly troubling for coral-specific studies and further indicate that care must be taken when comparing coral microbiome datasets resulting from even highly similar protocols.

Alpha diversity metrics on ASV data (both abundance-based and phylogenetic) were consistent between protocols in both coral species. Alpha diversity using OTU data were comparable between protocols with all metrics for *M. aequituberculata*, but significant differences were present with Shannon Diversity and Faith’s PD indices for *P. lobata*, suggesting some variability in alpha diversity in regard to both relative abundances and phylogenetic makeup when sequences are grouped with 97% similarity only. Nonetheless, our results suggest that comparisons of some alpha diversity metrics between protocols may be more reliable than comparisons of community composition. The results found here should be benchmarked in other systems and tested more broadly across species to determine the extent to which small differences in protocol might bias the perceived composition of host-associated or environmental microbiome sequencing.

Regardless of rarefaction, removal of low abundance reads, or comparisons of the resulting data at higher taxonomic levels, the bacterial community composition and relative abundances of taxa maintained differences between the two protocols but in ways that were inconsistent across host species and analytical metric. This can result, for instance, from one set of samples containing taxa that may never be present if they are prepared with a different protocol or sequenced on a different platform, likely due to differences in sequencing depth, where the deeper sequencing of the HiSeq platform can provide a greater opportunity to identify rare taxa [5], thus shifting the overall community composition. However, differences were also apparent between OTU and ASV datasets, suggesting that how we characterize bacterial species and/or strains, and at what taxonomic level we choose to analyse these data, may result in unintended biases. We found no evidence of differences in secondary structure of two differentially abundant taxa (*Vibrio* and *Endozoicomonas*) that could have resulted from differential denaturation of sequences in the two platforms due to differences in platform chemistry [38]. Specifically, there was no indication of high GC content in these sequences, which has previously been found to affect read numbers from Illumina sequencing runs due to intermittent halting of polymerase during amplification [39, 40]. While this was not an exhaustive dive into the effects of platform chemistry on sequencing outcome, it suggests that differential abundances of specific taxa are unlikely to be caused by the presence of differential secondary structures. However, further research is necessary to rule this out completely.

The samples used in this study were not initially intended to test differences between protocols or sequencing platforms, but rather provided an opportunity to examine an overlapping set of technical replicates that arose from a larger study comprised of similarly prepared and differentially sequenced samples. Thus, we cannot clearly identify the specific mechanism(s) involved in driving the found community differences. Biases in these complex microbial communities could be a result of 1) differences in sequencing depth that are not overcome by rarefaction or other *in silico* normalizations, 2) library denaturation and/or sequencing platform chemistry, 3) differences in the type of Hi Fidelity Taq used in PCR and potentially even 4) user and/or facility bias [41, 42]. Regardless, the results shown here reveal not only the necessity to design a targeted study to examine procedural and mechanistic differences in sequencing protocols, but also the responsibility of researchers to proceed with extreme caution when combining and interpreting datasets that are generated from subtly and seemingly innocuously different methodologies.

### Conclusions

The present study found limitations in our ability to compare coral microbiome ‘technical’ replicates that were generated in almost identical fashions but then sequenced on different platforms. Despite attempts to rectify these issues with some commonly used normalization methods, we still found significant differences in some alpha diversity metrics and in most beta diversity metrics between the two protocols. These inconsistencies make it difficult to identify a “cure-all” adjustment for comparability between even highly similar protocols and, instead, differences among protocols and sequencing platforms are more likely to be specific to the microbiome host and specific set of microbiomes found in each dataset. Studies that aim to compare beta diversity may find more confidence in their results if overlapping technical replicates for each dataset and host species are run to ensure correct adjustments are used for these specific datasets. Based on these results, we urge caution in the statistical comparison and interpretation of 16S rRNA datasets that combine data resulting from different protocols and sequencing platforms. While we continue to encourage meta-analyses to discover of cosmopolitan patterns in microbiome dynamics, we advise researchers to be cognizant that even minor variations in the protocol can significantly affect microbiome composition, and those running longitudinal studies be rigorous in the consistency of their methods through time.

## Acknowledgements

The authors would like to thank the Kiritimati Project Manager, K. Tietjen, for her contribution to the organisation and conduction of field work that provided samples for this comparative study, and to J. McDevitt-Irwin for personal communications about handling of the sequencing data. Thanks also go to the Earth Microbiome Project (EMP) and the Global Coral Microbiome Project (GCMP) who provided the funding and resources for the HiSeq sequencing, and to G. Ackerman for her unyielding assistance in data access. H.E.E. acknowledges support from the National Science Foundation (NSF) through a Postdoctoral Research Fellowship in Biology (PRFB). S.S. acknowledges support from a National Sciences and Engineering Research Council Postdoctoral Research Fellowship (NSERC PDF). J.K.B. acknowledges support from a National Sciences Foundation RAPID Grant (OCE-1446402), an NSERC Discovery Grant, the Canadian Foundation for Innovation, the Packard Foundation, a Pew Fellowship, and the University of Victoria. R.V.T acknowledges support from NSF. The raw data from the Illumina MiSeq samples are publicly available from McDevitt-Irwin et al. [11] at Harvard DataVerse: https://doi.org/10.7910/DVN/3QZTT1. The raw data from the Illumina HiSeq samples are available on the NCBI Sequence Read Archive (SRA) under the BioProject accession PRJNA687031. All bioinformatics and statistical codes are freely accessible via github at github.com/hannaheps/comparison.

